# SKA: Split Kmer Analysis Toolkit for Bacterial Genomic Epidemiology

**DOI:** 10.1101/453142

**Authors:** S. R. Harris

## Abstract

Genome sequencing is revolutionising infectious disease epidemiology, providing a huge step forward in sensitivity and specificity over more traditional molecular typing techniques. However, the complexity of genome data often means that its analysis and interpretation requires high-performance compute infrastructure and dedicated bioinformatics support. Furthermore, current methods have limitations that can differ between analyses and are often opaque to the user, and their reliance on multiple external dependencies makes reproducibility difficult. Here I introduce SKA, a toolkit for analysis of genome sequence data from closely-related, small, haploid genomes. SKA uses split kmers to rapidly identify variation between genome sequences, making it possible to analyse hundreds of genomes on a standard home computer. Tests on publicly available simulated and real-life data show that SKA is both faster and more efficient than the gold standard methods used today while retaining similar levels of accuracy for epidemiological purposes. SKA can take raw read data or genome assemblies as input and calculate pairwise distances, create single linkage clusters and align genomes to a reference genome or using a reference-free approach. SKA requires few decisions to be made by the user, which, along with its computational efficiency, allows genome analysis to become accessible to those with only basic bioinformatics training. The limitations of SKA are also far more transparent than for current approaches, and future improvements to mitigate these limitations are possible. Overall, SKA is a powerful addition to the armoury of the genomic epidemiologist. SKA source code is available from Github (https://github.com/simonrharris/SKA).

## Introduction

Genome sequencing of bacterial pathogens is rapidly becoming an essential tool in the epidemiologist’s armoury. It provides increased specificity and sensitivity over more traditional molecular typing approaches such as pulsed-field gel electrophoresis and multi locus sequence typing (MLST), as well as providing other epidemiologically-relevant data such as genotypic anitimicrobial resistance prediction. However, these improvements come at the cost of analytical complexity that often requires dedicated bioinformatics support to analyse genomic data and interpret results. Automation is possible, but typically requires complex analytical pipelines and significant compute infrastructure and some input from an experienced user is still often necessary. Here I present SKA (<u>S</u>plit <u>K</u>mer <u>A</u>nalysis), a toolkit that provides a means for rapid analysis of bacterial genomic data for epidemiological purposes. SKA is fast, memory efficient and produces only relatively small intermediate files, making it feasible to analyse hundreds of bacterial (or other small, haploid) genome sequences on an average home computer. SKA is particularly powerful for analysing closely-related genomes, such as those from an outbreak or transmission chain. It is not designed for population genomics applications at a species-wide or larger scale.

To make use of genomic data for epidemiological purposes, the first step is to create hypotheses of homology between bases in the sequences of the samples of interest. This can be based on a single common reference sequence against which data from all new samples are mapped, or by using reference-free techniques. Mapping is a relatively simple approach for identifying variation. Raw sequencing data is aligned to a relevant reference genome, usually a complete, closed genome sequence, using one of an array of software options. Aligned reads are then filtered and consensus base calls calculated for each base of the reference genome. Although mapping algorithms employ creative methods to speed up the alignment process, such as by indexing sequences using hash tables or burrows wheeler transforms (1), this process can still be relatively costly in terms of compute resources, including central processing unit (CPU) usage time, memory usage and disk space to store large intermediate files. The most common reference-free approach to genomic epidemiology involves analysis of a set of conserved genes. This core gene approach requires genome assembly followed by homology assessment, and often also alignment, of alleles. Neither reference mapping nor core genome methods are ideally suited for genomic epidemiology. Reference mapping can lead to biased results in cases where the reference is distant from all or some of the isolates in the analysis. These a biases are difficult to predict, and differ between analyses of the same data and even between samples within a single analysis. Core genome methods, on the other hand, exclude large regions of the genome, so can lack sensitivity.

### Rapid genomics: The kmer revolution

The massively increasing number of genome sequences available for both eukaryotes and pathogenic bacterial species has necessitated a new generation of analytical methods. Many of these methods have leveraged the computational efficiency of exact matching by breaking genomic data down into small pieces of a known size (k), often called kmers. Kmer matching is extremely fast and scalable to thousands of genome comparisons. Examples of some of the more revolutionary applications are Mash (2), for rapidly estimating approximate Jaccard distances (3) between genomes, minimap (4), for aligning long sequencing reads to reference genomes extremely quickly and BIGSI (5), for searching for matches in graph representations of extremely large genome databases. To further increase analysis speed and reduce memory requirements, these methods use minimisers (6) and/or bloom filters (7). For closely-related, small genomes, however, such approximations are not necessary. All kmers can be stored and searched without adversely impacting speed and efficiency. Even so, a big challenge for kmer-based approaches is sequence variation. Counting kmers can identify similarity, but non-matching of kmers cannot distinguish variation in the sequence from absence of that sequence, and thus kmer methods for calculating distances are most applicable to divergent samples. Identifying relationships between samples within a single MLST sequence type, for example, is often inaccurate using kmer distances.

### Rapid identification of variation using split kmers

Here I introduce the concept of the split kmer, which is simply a pair of kmers in a DNA sequence that are separated by one or more bases. Split kmers allow the speed of exact matching to be utilized to identify regions of variation flanked by conserved sequences. A related approach, whereby kmers are first counted and then sorted before looking for variation nat the middle base has been used in the kSNP software (8). kSNP does this by post-processing results from the jellyfish kmer counting software (9), which limits its speed, efficiency and versatility. Split kmers also have the added potential over the kSNP approach of allowing the analysis of insertions, deletions and other structural variations in genomes.

## SKA: A toolkit for split kmer genomic epidemiology

SKA was developed to allow rapid, simple analysis of short-read genome sequence data from bacteria or other haploid organisms with small genomes. It is written in C++ with no dependencies other than a UNIX-like command-line interface, gnu make and a version of g++ compatible with C++11 for installation from the makefile. Source code and documentation are available on github at https://github.com/simonrharris/SKA and https://github.com/simonrharris/SKA/wiki respectively. The SKA toolkit includes methods extract split kmers from assemblies or directly from fastq format read sequences, compute pairwise distances, identify clusters, produce sequence alignments with or without a reference sequence and provide various comparison and summary statistics. SKA comprises a set of subcommands to carry out these tasks, which can all be accessed via the single ska command. SKA currently uses the simplest form of split kmers, that is two kmers separated by a single base, to allow rapid comparison and alignment of single nucleotide polymorphisms (SNPs) in small, conserved genomes, making it particularly suited for bacterial pathogen surveillance, outbreak investigations and transmission tracking. Here I show with published simulated and real-world outbreak data that SKA is both computationally efficient and accurate, providing another powerful tool for the growing field of genomic epidemiology.

### Creating split kmer files

SKA stores split kmers in a new file format known as skf (split kmer format) files, which can store sequence information for one or more samples. These files have the following structure. The first line reports the version of SKA that created the file. The second line contains a single integer representing the split kmer size used to create the split kmers in the file. The split kmer size is the size of each of the two kmers making up the split kmer. The third line is a whitespace separated list of the samples in the file. Subsequent lines are made up of a compressed representation of the split kmers for various subsets of the samples in the file. SKA can create split kmers from fasta and fastq files using three subcommands: ska fasta, ska alleles and ska fastq. Ska fasta and ska alleles require fasta format input and differ only in the way they interpret these files. For ska fasta, the split kmers in all fasta sequences in the input file(s) are treated as belonging to a single sample, as would be required for a set of contigs from an assembly or reference genome. For ska alleles, each fasta sequence is treated as a separate sample, as would be the case if the input file was a set of alleles of a locus. Ska fastq creates split kmers from one or more fastq files. As with ska fasta, all sequences in the input fastq file(s) are treated as belonging to a single sample. Importantly, ska fastq carries out a number of simple filtering steps to reduce noise in the split kmer set produced. These include breaking sequences at Ns or at bases with a base quality below a user-defined minimum (default=20) and only creating split kmers for the subsequences with qualities above this level, requiring a user-defined minimum coverage for each split kmer per input file (default=2), requiring a user-defined minimum total coverage for each split kmer (default=4), and a user-defined minimum minor allele frequency (default=0.2), with kmer alleles below this being discarded. The base quality filter is particularly efficient at reducing the number of noisy kmers, and can dramatically reduce the number of split kmers stored for a sample. Similar approaches could provide major efficiency improvements for many other kmer-based methods. The coverage and minor allele frequency cutoffs are important for reducing uncertainty in middle base calls caused by sequencing error, and are analogous to the sorts of filter that are often applied when calling variation from pileups of reads mapped against a reference genome. However, applying them at the kmer creation stage further improves the efficiency of downstream analysis.

### Merging skf files

SKA is designed to be used for genomic epidemiology, particularly for outbreak and transmission investigations. The similarity of samples in these cases creates large amounts of redundancy in split kmer sets between samples. This redundancy can be utilized to improve the efficiency of data storage and, as a consequence, data access by merging individual skf files using ska merge. Most SKA commands that take skf files as input can read merged files, and, given a file containing a list of samples, can analyse subsets of samples from merged skf files. Individual sample skf files can be re-extracted from merged skf files using the same approach.

### Summarising and comparing skf files

SKA provides two subcommands for summarising split kmer files. Ska info provides information about the content of split kmers files, including the split kmer size, number of samples and number of split kmers. The ska summary command produces a tab separated table reporting a number of useful quality control (QC) statistics for each sample in the file(s), including the split kmer size, the total number of split kmers, the number of middle bases that are As, Cs, Gs, Ts and Ns and the GC content of the middle bases. For QC purposes, if the species of the sample is known it would be expected that the total number of split kmers should be approximately the same as the genome length, while the GC content of the middle bases should be representative of the species. It should be noted that repeat regions will reduce the number of split kmers while sequencing error and contamination may increase the number. In cases where the species is unknown, the number of split kmers can be checked to make sure it is in a sensible range for a bacterial sample (e.g. usually within a range of around 1Mb to 10Mb).

Ska compare allows samples in a query skf file to be compared with those in one or more other skf files. The pairwise output includes the number of split kmers unique to each sample, the number of matching split kmers and the number of SNPs identified between the samples.

### Pairwise distance calculation and single linkage clustering

Ska distance calculates a number of pairwise indices and distances between samples as well as performing single-linkage clustering of samples based on a user-defined SNP cutoff (default=20). The statistics reported for each pair of samples are shown in table 1.

**Table 1.** Description of the columns in the ․distances.tsv output of ska distance.

| Column | Description |
| --- | --- |
| **File 1** | The name of the first sample being compared |
| **File 2** | The name of the second sample being compared |
| **Matches** | Number of split kmers found in both samples where the middle base is an A, C, G or T and matches between samples |
| **Mismatches** | Number of split kmers found in only one of the samples |
| **Jaccard Index** | Ratio of split kmers found in both samples to the total found in the two samples: $\text{matches}/(\text{matches}+\text{mismatches})$ |
| **Mash-like distance** | A distance based on the Mash distance calculation using the Jaccard Index (j) above and the split kmer length (k): $(-1/(2k+1)) * \ln(2j/(1+j))$ for $0 < j \leq 1$ or 1 for $j=0$ |
| **SNPs** | Number of split kmers found in both samples where the middle base is an A, C, G or T but differs between files |
| **SNP distance** | The ratio of SNPs to matches: $\text{SNPs}/\text{matches}$ |

Clusters are reported in a tab delimited file, and the sample names in each cluster containing multiple samples are output to individual text files that can be used to subset further analyses. Finally, a dot format file representing the clusters is produced to facilitate visualisation, for example in microreact (10).

### Aligning split kmers

One of the key features of SKA is the ability to create reference-free genome alignments with ska align. This is simply a kmer matching exercise, with the middle bases of matching split kmers output as the alignment. It is important to note that there is no positional information in the output alignments, which are therefore not suitable for analysis in recombination detection methods, such as Gubbins or ClonalFrameML, that utilise SNP density. Output alignments can be filtered to only include split kmer alignments that were present in a user-defined proportion of samples (default=0.9). There is also the option to limit the output alignment to variable sites, in which case the number of constant sites of each base, where all aligned middle bases are the same base or N, are output to screen. Some phylogenetic methods can use this information to correct for ascertainment biases inherent in variable site alignments.

### Mapping split kmers to a reference

Ska map allows split kmers to be aligned to (or mapped) against a reference genome. By default, the output alignment will only include sites mapped to by the middle base of a split kmer. However, there is also the option to map all bases covered by a split kmer. This improves coverage and allows mapping in the bases surrounding a variant site. As with ska align, the output can be restricted to variant sites only, in which case the number of constant sites for each base are output to screen. Due to the necessity for exact matches of split kmers, ska map works best when the reference is very similar to the samples being mapped (see limitations section below).

### Annotating split kmers

Kmer-based analysis can sometimes seem separated from biological reality, with no obvious way to relate variation to a genomic context. For the reference-free methods of SKA that is particularly true. SKA includes the annotate subcommand to allow split kmers to be located in genomes, and where annotation is present, to extract biologically relevant information about the variant. Ska annotate accepts fasta and gff reference input and outputs a vcf. The vcf reports the chromosome and position of all middle base mappings to the reference sequence along with the reference and alternate bases along with various information in the info field (see Table 2). As with other SKA subcommands, the output can be restricted to variant sites only.

**Table 2.** Description of values in ska annotate vcf output file info column. Gff only indicates where values are only output when the input is a gff file. Optional indicates where a value is only output when requested by the user.

| ID | Type | Description | gff only | Optional |
| --- | --- | --- | --- | --- |
| NS | Integer | Number of samples with a matching split kmer |  |  |
| NS5 | 5 Integers | Number of samples with A, C, G, T or N as the middle base |  |  |
| FT | String | Feature type | X |  |
| FS | Character | Feature strand (+/-) | X |  |
| BP | Integer | Position of base in feature | X |  |
| CP | Integer | Position of base in codon | X |  |
| AAP | Integer | Position of amino acid in feature | X |  |
| RAA | Character | Reference amino acid | X |  |
| AAA | Character(s) | Alternate amino acid(s) | X |  |
| ID | String | Feature ID | X |  |
| LT | String | Feature locus tag | X |  |
| SI | String | Feature systematic ID | X |  |
| GE | String | Gene | X |  |
| PR | String | Product | X | X |

### Weeding split kmers

When calling variation between genomes, particularly for the purpose of phylogenetic reconstruction, it is often useful to exclude parts of the genome that are known to be transmitted horizontally rather than vertically. Traditionally this can be achieved by masking regions of a reference genome during read mapping, or by analyzing core genome gene sets. Typically, kmer-based analyses do not exclude these accessory regions. Ska weed allows kmers matching known accessory sequences to be excluded from split kmer files. For example, to remove known phage, plasmid and other mobile sequences, the user needs to simply create a split kmer file from sequences of these elements and weed them from split kmer files of the samples of interest using ska weed. Other known contaminants can be excluded in the same way.

### Identifying unique split kmers

Ska unique outputs a set of kmers that are found in an ingroup set of samples but not found in any other samples. For outbreak analyses this allows the creation of outbreak specific split kmer sets which can be used to rapidly screen new isolates to see if they are members of the outbreak using ska compare, which outputs various comparison statistics between a query skf file and one or more others. This technique works best if relatively close outgroup isolates are included.

### Typing alleles using split kmers

SKA includes a typing subcommand, ska type, which accepts a set of multifasta files for alleles of loci of interest and an optional profiles file which translates allele numbers to sequence types. This can be used, for example for rapid MLST from skf files. Ska type will export sequences of novel or partial alleles, making it more adaptable than some other kmer-based sequence typing methods. It should be noted, however, that ska type does not handle diverse alleles and repetitive alleles well due to the limitations imposed by the use of split kmers (see limitations section below).

### Limitations

The use of kmers in general, or more specifically split kmers comes with a number of limitations. In particular, split kmers lose power as the density of variants increases. This is simply because split kmers no longer match if a second variant site occurs within a kmer length of either side of a middle base. For this reason, split kmers are particularly suited for the analysis of highly similar sequences, although results for analyses of more diverse sequences are still good in many cases (e.g. see results section below). This limitation affects different analyses slightly differently. For pairwise methods, such as pairwise distance calculation or mapping to a reference, the diversity between each pair will impact the results. For these analyses SKA outputs an estimate of the number of alignments that have been missed due to this phenomenon. This is purely a guide and should not be considered an accurate count of missed alignments. For reference-free alignment of split kmers the situation is more complex, since in an all versus all alignment the total number of variant sites in all samples becomes important. This is because variants between any samples that are within a kmer length of each other will lead to missed alignments. Therefore, ska align outputs a prediction of the total number of missed alignments in the entire set of samples being aligned. In practice these often occur on longer branches of the tree and lead to subalignments within the data, which can be accessed by changing the minimum proportion of samples required to output an alignment. In cases where large numbers of alignments are missed, reanalyses of subsets of closely related samples (e.g. by subclade) can be used to improve resolution of those clades.

A second limitation of using split kmers is that repeats of the kmer size or greater cannot be analysed, even if the raw sequence reads would include unique sequence either side of the repeat. This is simply because information linking split kmers to others in the same reads are not stored. In practice this usually results in a minor loss of power.

A third consideration when using split kmers is that variation in the first or last k bases of the sequence (where k is the kmer length) cannot be identified for linear input sequences. For fasta files of reference genomes, ska fasta and ska map allow the sequence to be circularized to avoid this problem. For fasta input of uncircularised contigs or fastq files this is not possible. However, the first and last few bases of assembled contigs and sequencing reads are known to be the least accurate, so this may have the effect of reducing some noise in the data. It does, however, impact on depth of coverage for fastq file input.

In general, split kmer analysis requires relatively high-quality input sequence data, for example from Illumina technologies or polished assemblies. Poor quality sequencing with high error rates or low coverage can lead to poor results. At the time of writing raw reads from long read technologies are not of a high enough quality for use in SKA. Reduction of the ska fastq filtering options can rescue data from poor sequencing runs, but often at the expense of increased noise which may impact on speed, memory usage and accuracy of analyses. Therefore, it is recommended that long read data is assembled and corrected before using with SKA.

Finally, the current version of SKA only supports split kmers with a single base in the middle. Split kmer analyses could extend upon this to allow analyses of more complex data, including indels and short repeats.

## Methods

Unless specifically mentioned, SKA commands were run using default options in SKA v1.0.

### Analysis of simulated data from Lees *et al.* (11) Methods

The ability of SKA to correctly reconstruct the phylogeny from the simulated genomic data was assessed using three approaches:

1. Calculation of a pairwise distance matrix using SKA distance
2. Alignment of split kmers against a reference using SKA map
3. Reference-free alignment of split kmers using SKA align

The raw simulated read data and assemblies and inferred trees of Lees *et al.* (11) were retrieved from figshare at https://dx.doi.org/10.6084/m9.figshare.5483461 and https://dx.doi.org/10.6084/m9.figshare.5483464 respectively. Split kmers were created from sample fastq files using ska fastq and from assemblies using ska fasta. Ska map was run using the TIGR4 genome (12) as reference with default options and with the option to only output variant sites. Ska align was run with the minimum proportion of isolates required to possess a split kmer for that kmer to be included in the alignment set to zero given the diverse nature of the data. Ska distance was run with default options to calculate pairwise distances.

Trees based on the distances from ska distance were reconstructed using bioNJ calculated in R v3.5.1. Distances were first rescaled using the rescale function from the scales package v1.0.0, and trees were calculated using the bionj function in the Ape package v5.1 (13). RapidNJ v2.3.2 (14) trees were run on ska map and ska align alignments under default options (Kimura model of evolution). Maximum likelihood trees were reconstructed from ska map and ska align alignments using IQ-Tree v1.6.5 (15) with and without the fast option. For all analyses a general time reversible (GTR) (16) model of evolution with a discrete gamma model (17) with four rate categories for among site rate variation was used. For analyses of alignments of variant sites only, an ascertainment bias correction (18) was applied. kSNP3 v3.1 was run from the assemblies of Lees *et al.* using a kmer length of 15 to give consistency with the kmer size used in SKA. All other options were set as the default. Trees were run from the resulting alignment using IQ-Tree with the fast option and a general time reversible (GTR) model of evolution with a discrete gamma model with four rate categories for among site rate variation.

Kendall Colijn metrics (19) were calculated between kSNP3 trees, SKA trees and trees from Lees *et al.* with the real tree from Lees *et al.* in R v3.5.1 using the treespace package v1.1.3 (20) with Ape v5.1. Trees were first midpoint rooted using the midpoint function from the Phangorn package v2.4.0 (21), followed by running treespace with an nf of 3 and a lambda of 0. Robinson Foulds distances (22) were calculated in python v2.7.13 using the symmetric_difference function of dendropy v4.2.0 (23).

### Analysis of outbreak datasets from Timme *et al.* (24)

For all datasets split kmers were created from raw reads of each sample, where available, using SKA fastq with default options. For the *E. coli* dataset, split kmers were produced for sample 2011C-3609 using ska fasta from the reference genome sequence. Ska distance was used with default options to create single-linkage clusters of samples separated by fewer than 20 SNPs and ska align used to create an alignment of variant sites with all other parameters set to the default. Based on the results of the analyses of the simulated data from Lees *et al.* phylogenetic trees were reconstructed from these alignments using IQ-Tree v1.6.5 with a GTR+G+ASC model and the fast option. The phylogenetic trees and the cluster networks dot files produced by ska distance were visualized in Microreact (10) along with a csv combining metadata for the samples with the ska distance clusters.

### Analysis of MRSA outbreak data from Harris *et al.* (25) and Coll *et al.* (26)

Split kmers were created for the 65 samples from Harris *et al.* using ska fastq with default options before being merged using ska merge. QC statistics were produced with ska summary and pairwise distances and single linkage clustering performed with ska distance with default parameters (SNP cutoff (-s) 20, minimum identity (-i) 0.9). Split kmers unique to the 45 outbreak samples were identified using ska unique, by providing it with the file containing a list of the 45 sample names created by ska distance as one of its output cluster files.

To illustrate how SKA can be used to identify novel cases linked to an outbreak, the sequence data of the 1,683 samples from *Coll et al*. that had been typed by them as being part of MLST clonal complex 22, were downloaded from the European Nucleotide Archive and ska fastq and ska merge used to create a merged skf file from these samples. QC statistics for this merged skf file were produced with ska summary. The split kmers unique to the 45 outbreak samples from Harris *et al.* were compared with the merged split kmers from the samples of Coll *et al.* using ska compare. Results were sorted using UNIX sort with the ‐g and ‐r flags and the ‐k option set to 5 to sort by the percentage of split kmers in each sample matching the query split kmers. The skf files from the 45 outbreak samples from Harris *et al.* and the eleven samples with more than 5% match to these from the data of Coll *et al.* were then merged using ska merge. Ska distance was run on this merged skf file under default options. To compare the ability of SKA to reconstruct phylogenetic relationships between outbreak samples, the skf files from the 45 outbreak samples from Harris *et al.* and the ten samples from Coll *et al.* that were linked to the outbreak were merged using ska merge. A reference-free alignment was produced from this merged skf file using ska align with the minimum proportion of isolates required to possess a split kmer for that split kmer to be include in the alignment set to 0.9 (default), the ‐v flag set to output a variant only alignment, and the ‐k flag set to output the split kmers of variant sites to a file. This skf file of variant sites was used to identify the location of the bases in the reference-free alignment in the HO 5096 0412 and GCA_001369415.1 reference genomes using ska annotate to allow comparison of the SNPs identified by SKA with those from the output of SNP Pipeline v2.0.2 (27). SNP pipeline was run using default parameters against two reference genomes: HO 5096 0412 as an example of a realistic reference and GCA_001369415.1, a complete genome of one of the outbreak samples from Harris *et al.*, as an unrealistically close reference. In both cases, the individual steps of the pipeline were run manually to allow computational resources to be accurately recorded. SNP pipeline has a number of dependencies. The versions of these used in the analysis were: Bowtie2 v2.2.3 (28), SAMtools v1.6 and BcfTools v1.5 (29), Picard v 2.18.14 (30), GATK v3.4-46 (31), VarScan v2.3.9 (32) and tabix v1.8 (33). Mapping was done using Bowtie2, and maximum memory allocated to GATK and Picard was restricted to 5Gb. All other parameters were set to the default. Any sites in the resulting alignments that failed filters or were constant within the outbreak samples (i.e. only differed in the reference) were removed. Phylogenetic reconstruction of the alignments from SKA and SNP Pipeline were carried out with IQ-Tree v1.6.5 with a GTR+G+ASC model and the fast option. SNPs were reconstructed onto the branches of the trees using pyjar (34), a python implementation of the joint ancestral reconstruction method of Pupko *et al.* (35).

## Results

To test the utility of SKA, it was applied to 7 published datasets from four publications:

1. A simulated dataset designed to compare methods for genomic phylogenetics (11).
2. Four benchmarking outbreak datasets for genomic epidemiology, comprising 23 *Campylobacter jejuni*, 9 *Escherichia coli*, 31 *Listeria monocytogenes* and 22 *Salmonella enterica subsp. enterica* ser. Bareilly (24).
3. 65 *Staphylococcus aureus* samples from an investigation into an outbreak on a special care baby unit (SCBU) at Addenbrooke’s Hospital, of which 45 have been previously identified as a single outbreak (25).
4. 1,683 ST22 samples from a genomic survey of *S. aureus* from the East of England (26).

All timings were made on an AMD Opteron (TM) Processor 6272 2.1GHz.

## Simulated data

### Description

Lees *et al.*(11) produced a simulated dataset for comparison of methods for genomic phylogenetic reconstruction. Sequences were simulated on the tree topology from a published analysis of a *Listeria monocytogenes* dataset (36) and the evolutionary model parameterised using values from *Streptococcus pneumoniae*. They provided raw sequence reads for 96 simulated genomes along with assemblies produced from those reads using a typical assembly approach. After analyzing the data with various phylogenetic approaches they compared the trees produced by each method with the true tree using the Kendall Colijn (KC) metric (19). They found methods using maximum likelihood phylogenetic reconstruction based on mapping raw reads against a close reference were the most accurate. These methods were also computationally expensive. The most accurate reference-free approach was bioNJ reconstruction using distances calculated with Mash (2). Although this method was considerably less accurate, it was one of the most computationally efficient approaches in the tests.

### Results

Supplementary Table 1 shows the KC and Robinson Foulds (RF) distances (22) to the true tree for each of the trees created using SKA along with trees from some of the better performing methods assessed by Lees *et al.* and kSNP3 as a comparison of an approach that is similar to SKA. Selected results are summarized in Fig. 1.

**Figure 1.**
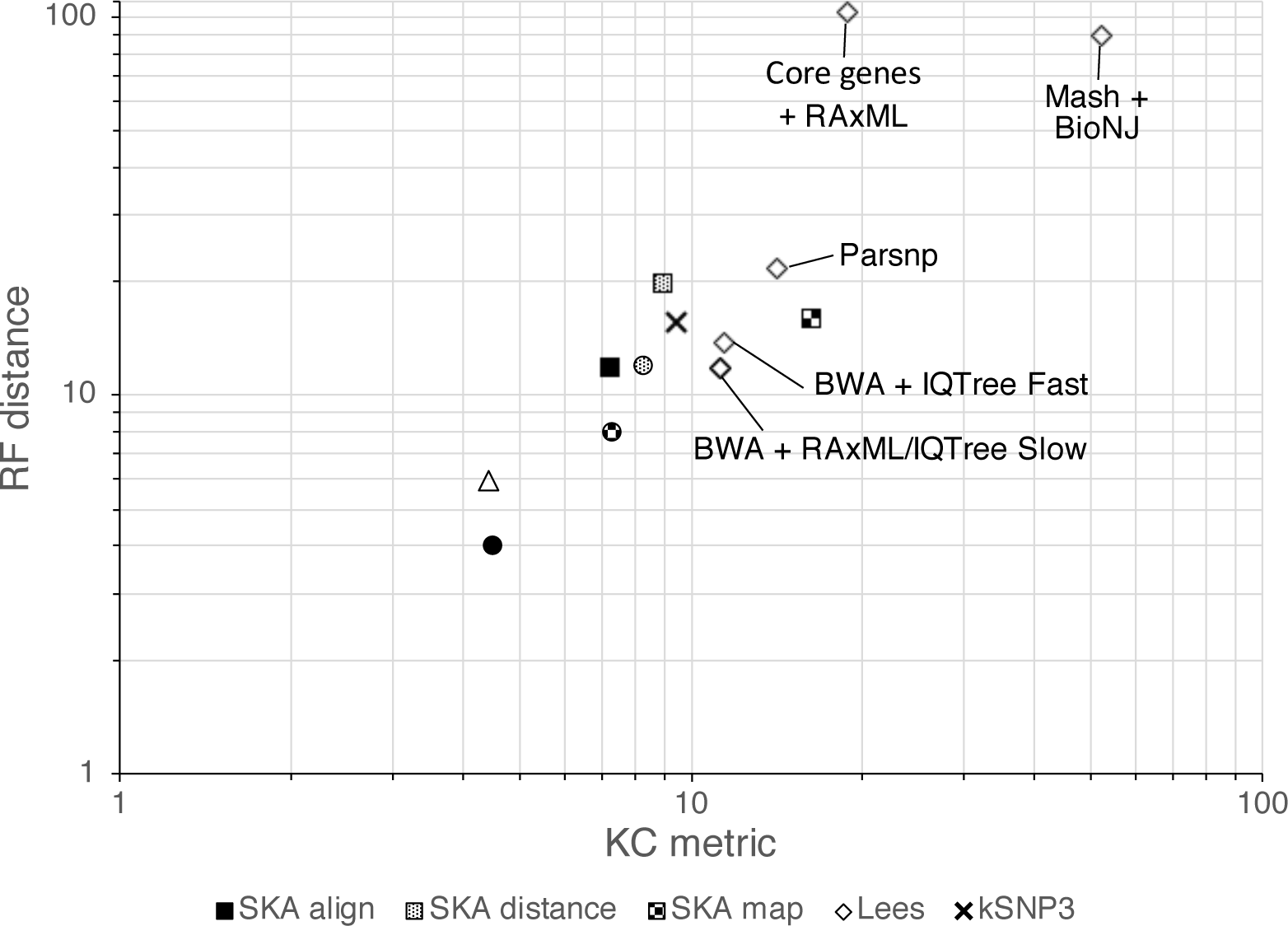
Comparison between SKA and other methods for their accuracy of reconstruction of the true tree from simulated read data of Lees *et al.* Kendall Colijn (KC) metric distances from the true tree are shown on the x axis, and Robinson Foulds (RF) distances from the true tree on the y axis. The key shows the fill patterns for trees reconstructed with different SKA commands. Trees from the analyses of Lees *et al.* are filled white. For SKA analyses, IQ-Tree fast trees are shown for alignments of variant sites from ska map and ska align, while BioNJ trees are shown for ska distance mash-like distances. Square markers indicate SKA analyses from assemblies and circles from raw sequencing reads. The black cross indicates the tree from kSNP3. The white triangle indicates the best performing tree from Lees *et al*, which was a RAxML tree of a BWA mapping to an artificially close reference genome. White diamonds represent trees from other analyses run in Lees *et al.* (11), and are individually labelled.

SKA preformed very competitively, with results from mapping, reference-free alignment and distance-based trees comparable to the best mapping approaches presented by Lees *et al.* Results based on split kmers from assemblies and raw reads were similar, although in general analyses from reads proved slightly more accurate.

The best performance resulted from reference-free alignments from SKA align. IQ-Tree reconstructions from these alignments produced trees of similar quality to the artificial gold standard method of Lees *et al.*, which used RAxML on a BWA alignment to an artificially close reference. RapidNJ trees were only marginally inferior when run on the variant sites only. A general theme of the results was that RapidNJ provided less accurate trees when run on all sites in any alignment. The KC metric was particularly poor for these trees, while the RF distance was only slightly worse than that for the RapidNJ trees from alignments of variant sites. Since the KC metric is particularly harsh on deep rearrangements, this suggests that RapidNJ trees from alignments of all sites failed to accurately reconstruct the deeper relationships in the tree.

Similar patterns of results were observed for alignments from SKA map, although the resulting trees were less accurate in all cases, and less accurate than the RAxML and IQ-Tree reconstructions from BWA mapping reported in Lees *et al.* This is unsurprising given the high level of variation of many samples from the reference genome, and the limitations of split kmers in those situations (see limitations section above).

BioNJ trees based on SKA mash-like distances, provided a similar level of accuracy to those of RapidNJ run on the reference-free alignments of variant sites from SKA align, and were more accurate than any of the realistic methods presented by Lees *et al.* Trees created from SKA SNP distances were similar to those of RapidNJ run on the reference-free alignments of all sites from SKA align, with the same loss of accuracy under the KC metric. kSNP3, which uses a similar approach to SKA also performed well in terms of the tree it produced, providing results comparable to mapping approaches and SKA. However, kSNP3 can only be run from assemblies and its runtime was much longer, memory requirements were higher, and disk space requirements far greater than SKA (see Supplementary Table 2). At one point during the analysis, kSNP3 had produced in excess of 50,000 intermediate files requiring more than 15Gb disk space.

## Outbreak benchmark datasets

### Description

Timme *et al.* (24) produced a public resource for benchmarking phylogenomic methods. The resource comprises four real foodborne pathogen outbreak datasets for which sequence data, gold-standard trees and epidemiological data are all available. The gold-standard trees were created using SNP Pipeline (27), a reference-mapping approach used to create an alignment, and Garli (37) for phylogenetic reconstruction. To assess the ability of SKA to identify outbreaks from real data without the need of a reference sequence, we analysed the four real outbreak datasets from Timme *et al.* with SKA.

### Results

Table 3 details the CPU time taken for each step of these analyses. In all cases, analyses took well under two hours of CPU time on a single core from raw data. The vast majority of this time was taken up creating split kmers from read data. Production of clusters, alignments and phylogenetic reconstructions from these split kmer files required less that 1.5 minutes total CPU time for all datasets. Clustering and phylogenetic results for the four datasets are shown in Fig. 2 and are available as interactive microreact instances at https://microreact.org/project/SKACjejuni, https://microreact.org/project/SKAEcoli, https://microreact.org/project/SKALmonocytogenes and https://microreact.org/project/SKASenterica.

**Figure 2.**
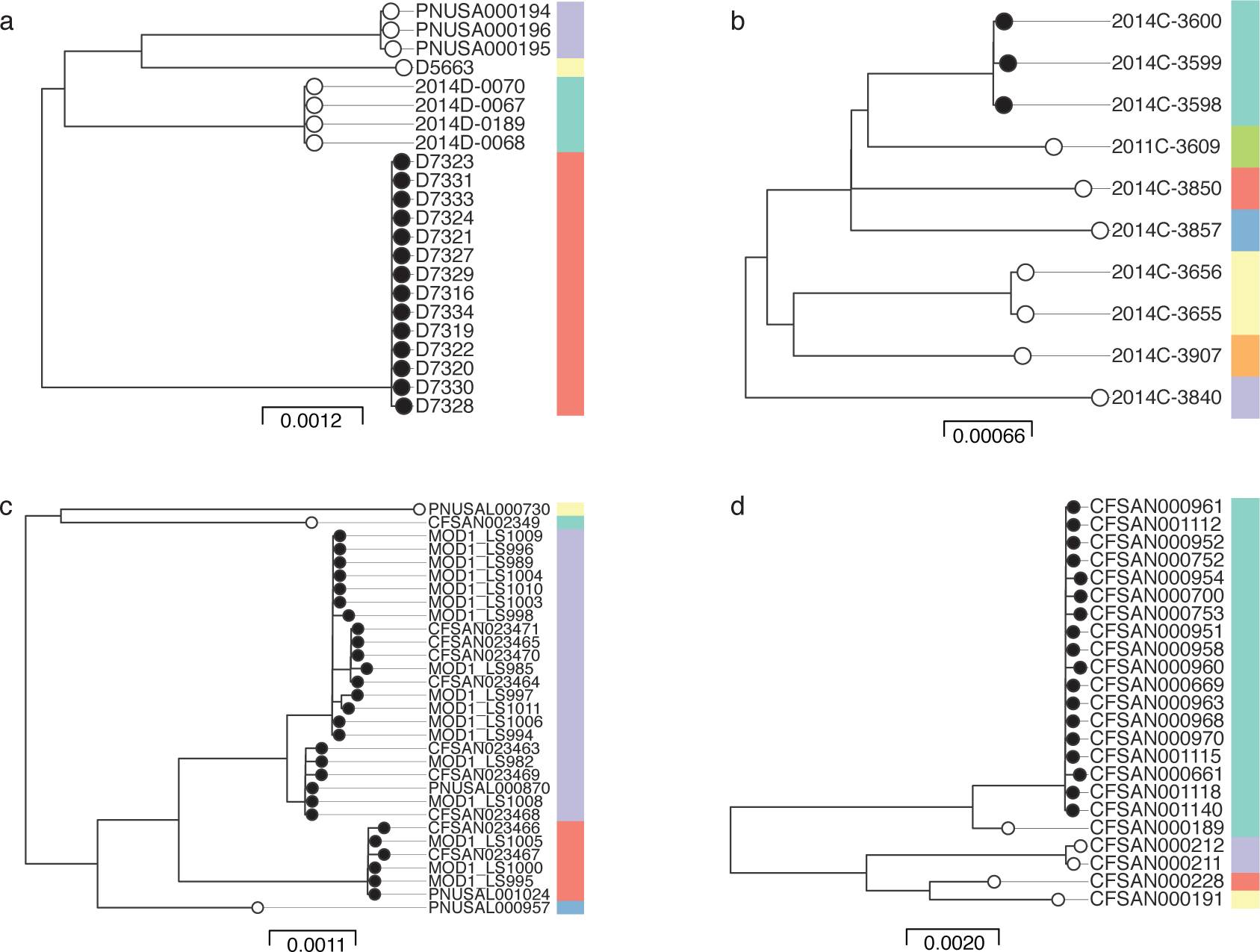
Results of SKA analysis four datasets from Timme *et al.* (24) a) *C. jejuni*. b) *E. coli*. c) *L. monocytogenes*. d) *S. enterica*. In all cases, trees shown were reconstructed from ska align alignments of variant sites only using IQ-Tree with a GTR+G+ASC evolutionary model and the fast option. Black circles on terminal nodes indicate isolates that were part of an outbreak according to the original investigation. White circles on terminal nodes indicate outgroup isolate that were not considered part of the outbreak. Coloured bars to the right of each tree indicate single linkage clusters identified by ska distance using a 20 SNP cutoff. Scale bars indicate substitutions per site on branches as output by IQ-Tree. All figures edited from microreact output.

**Table 3.** Data summary and CPU time for each stage of the SKA analyses of the four datasets from Timme *et al.* (24)

| **Dataset** | ***C. jejuni*** | ***E. coli*** | ***L. monocytogenes*** | ***S. enterica*** |
| --- | --- | --- | --- | --- |
| **Number of fastq files** | 22 | 9 | 31 | 23 |
| **Mean number of fastq reads** | 3,361,254 | 3,361,254 | 1,777,006 | 1,859,063 |
| **Mean number of fastq bases** | 657,706,314 | 356,661,903 | 380,854,631 | 278,574,418 |
| **SKA fastq mean CPU time (s)** | 271.35 | 145.21 | 184.29 | 118.14 |
| **SKA fastq total CPU time (s)** | 5,969.7 | 1,306.89 | 5,712.99 | 2,717.22 |
| **Number of fasta files** | 0 | 1 | 0 | 0 |
| **Mean number of fasta contigs** | - | 7 | - | - |
| **Mean number of fasta bases** | - | 5,412,686 | - | - |
| **SKA fasta mean CPU time (s)** | - | 10.06 | - | - |
| **SKA merge CPU time (s)** | 29.96 | 41.02 | 44.34 | 54.39 |
| **SKA distance CPU time (s)** | 8.22 | 13.24 | 8.51 | 15.71 |
| **SKA align CPU time (s)** | 10.29 | 14.06 | 12.59 | 19.24 |
| **IQ-Tree fast CPU time (s)** | 0.12 | 0.31 | 0.49 | 0.42 |
| **Total CPU time (s)** | 6,018.29 | 1,385.58 | 5,778.92 | 2,806.98 |
| **Total CPU time (m)** | 100.31 | 23.09 | 96.32 | 46.78 |
| **Total CPU time from split kmers (s)** | 48.59 | 68.63 | 65.93 | 89.76 |

In all four cases the phylogenies produced by IQ-Tree were similar to the trees reported by Timme *et al.* and recapitulated the expected clusterings. The SKA distance clusterings identified the expected outbreak sample sets for the *C. jejuni* and *E. coli* datasets. For the *L. monocytogenes* dataset, SKA split the outbreak into two clusters, which is not surprising given the clear diversity between those two clusters and is consistent with the IQ-Tree phylogenetic reconstruction and the trees reported by Timme *et al.* For the *S. enterica* dataset, SKA included one outgroup sample, CFSAN000189, in the outbreak cluster, because it was fewer than 20 SNPs from some isolates in the outbreak. However, the phylogenetic reconstruction clearly separated this outgroup from the outbreak isolates. In all four datasets SKA distance identified other, smaller clusters that were not the focus of the original outbreak investigations. These results highlight the problematic nature of using SNP cutoffs for outbreak definition, but illustrate that SKA provides rapid, accurate results that can be used to inform outbreak investigations without the need of a reference genome sequence.

## Case Study: MRSA outbreak investigation

### Description

In 2011 three babies in a special care baby unit (SCBU) at Addenbrooke’s Hospital in Cambridge, UK simultaneously tested positive for methicillin resistant *S. aureus* (MRSA) with near-identical Vitek antimicrobial resistance phenotypes during routine screening. The hospital deep cleaned the ward and launched an outbreak investigation. They found that over the previous six months 12 screening swabs had tested positive for MRSA with the same resistance pattern. However, there were two periods of time during those six months where no carriage was detected on the ward, making it difficult to conclude whether or not all of the cases were linked. Whole genome sequencing on the Illumina MiSeq platform was used by Harris *et al.* (25) to attempt to resolve this uncertainty. Twenty-six isolates were sequenced, including 17 isolates from the SCBU plus 9 isolates from other parts of the hospital that had no more than one difference in Vitek antibiograms from the outbreak samples. The results showed that 14 cases from the SCBU were linked, of which two were missed by the phenotypic method due to erroneous Vitek results. Further sequencing of community samples showed that the outbreak had spread into the community via the families of babies on the wards. During the investigation, two months after the last case, another baby on the ward tested positive for MRSA. This sample was immediately sequenced and confirmed to be part of the outbreak. Sampling of staff identified an individual carrying an MRSA that was confirmed to be linked to the outbreak by genome seuqencing. Decolonisation of the baby and staff member resolved the outbreak. In all, the genomic analysis linked 45 isolates to the outbreak and excluded a further 20.

The genomic investigation of the outbreak was carried out using a mapping approach, whereby sequenced reads were aligned to the closest available complete reference genome (HO 5096 0412) to identify variation between samples. Here, the data generated during the investigation was used to illustrate how SKA can be applied in such outbreak investigations and how it can quickly identify new linked samples. The CPU time and memory usage for SKA commands run during these analyses are shown in Supplementary Table 3.

### Creation and QC of split kmer files

For each of the 65 samples analysed in the outbreak investigation, ska fastq was run on the paired fastq files of raw reads using the default filtering options (minimum base quality of 20, minimum coverage per split kmer of 4 and minimum coverage per split kmer per file of 2). Ska summary showed that the split kmer files contained between 2,097,419 and 2,841,127 (mean=2,761,345, median=2,780,919) split kmers with GC contents between 31.7% and 32.7% (mean=32.6 %, median=32.6%). The reference ST22 genome HO 5096 0412 is 2,832,299 bases long with a GC content of 32.8%, so these numbers are consistent with expectations. Two samples had fewer than 2,500,000 split kmers, suggesting they were from poor sequencing runs where sequencing depth and/or quality dropped below the levels of the default ska fastq filters for parts of the genome.

### Pairwise distance calculation and clustering

Running ska distance with default options (SNP cutoff (-s) 20, minimum identity (-i) 0.9) clustered the 65 samples into 16 clusters, of which four contained two or more isolates. The largest cluster contained 45 isolates and corresponded to the outbreak samples identified in the original investigation. Two other clusters each contained two samples, and one contained four samples. In each case these were not isolated on the SCBU and potentially represent other outbreaks or transmissions that were not the focus of the published investigation. The maximum distance between any of the 45 isolates from the SCBU outbreak was 12 (minimum=0). Therefore, using SKA the SCBU outbreak could have been identified directly from raw sequencing reads in under two hours on a single CPU without the need for any prior information to identify a relevant reference genome, or even to know the species involved (total CPU time for all ska fastq, ska merge, ska summary and ska distance commands: 105.4 minutes. Maximum memory usage: 1.4Gb; Supplementary Table 3).

### Identifying new linked cases

SKA introduces a novel approach for efficient identification of new cases of a known outbreak. The first step is to identify conserved and unique split kmers from the known outbreak isolates using ska unique. The closer the outgroup samples are to the outbreak the more specific the method becomes. Using the 45 SCBU outbreak samples as the ingroup and the remaining 20 samples that were not identified as part of the outbreak as the outgroup identified 1,714 split kmers that were unique to the outbreak isolates. As new samples are sequenced, ska compare can be used to compare these unique outbreak kmers with the split kmers files produced from the fastq files of the new samples. This is extremely efficient. Toleman *et al.* (38) described ten further MRSA cases in the East of England from the following year that were found to be linked to the SCBU outbreak based on genomics and epidemiology. These isolates were discovered in a genomic survey of MRSA in patients in the East of England over a 12-month period from April 2012 to April 2013 by Coll *et al.* (26). They sequenced 2,282 isolates from 1,465 patients and found that the most common MLST clonal complex (CC) was CC22, the same CC to which the SCBU outbreak samples belonged. The ten isolates identified by Toleman *et al.* were part of this survey and were initially identified as linked to the SCBU outbreak on the basis of their MLST profile matching the novel profile of the SCBU outbreak samples. To assess whether SKA could identify the samples linked to the SCBU outbreak from the East of England survey, read data for the 1,683 samples listed as belonging to CC22 in the supplementary data table of Coll *et al.* were downloaded. Split kmers were created for each pair of fastq files using ska fastq and then merged into a single file. Fig. 3(a) illustrates the compression gained by storing these data in a merged skf file. QC of the data using ska summary (Fig. 3b) shows that a number of samples produced more split kmers than expected, suggesting they may contain DNA from contaminant sources, which will have the effect of inflating the size of the merged skf.

**Figure 3.**
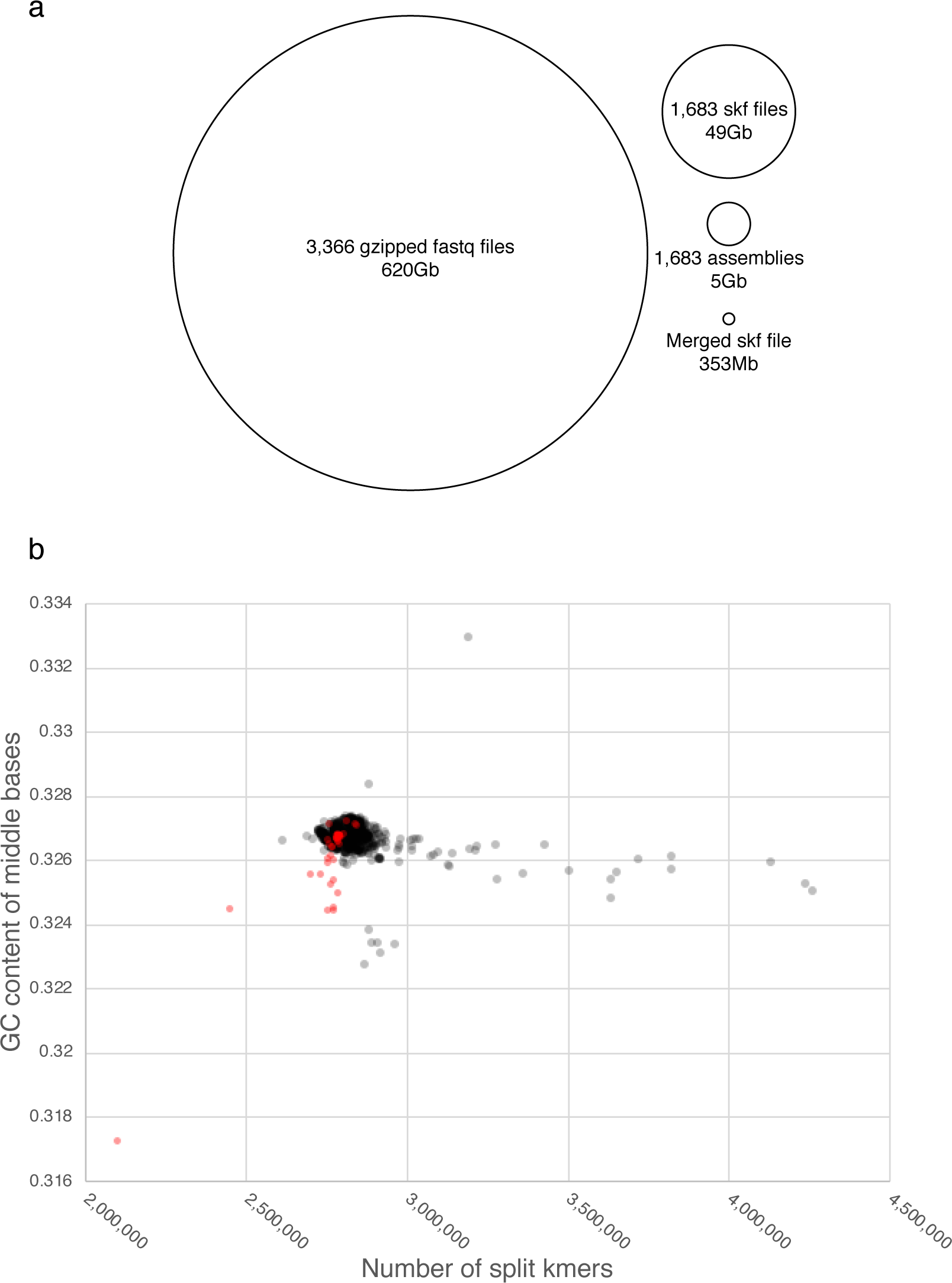
Summary and QC of split kmers created from the 1,683 CC22 MRSA samples of Coll *et al.* (26) and the 45 outbreak samples from Harris *et al.* a) Illustration of the data compression of a merged skf file containing all 1,683 samples relative to individual skf files, genome assemblies and gzipped fastq files. Areas of each circle are proportionate to the total size of the files. b) QC results from ska summary showing the number of split kmers and GC content of the middle bases of split kmers for the 1,683 MRSA samples from Coll *et al.* (black) and the 45 SCBU outbreak samples from Harris *et al* (red). Contaminated samples contain far more split kmers than expected for a genome of the length of *S. aureus*, while samples for which sequencing quality was poor contain fewer split kmers than expected.

Scanning the 1,683 samples from Coll *et al.* for the unique kmers of the SCBU outbreak samples required just 116s and 7Mb memory and identified ten samples matching 100% of the split kmers with no SNPs. One further isolate matched 34% of the split kmers with 34 SNPs. No other isolates matched more than 5% of the split kmers. To confirm whether the 11 isolates with greater than 5% match to the unique outbreak split kmers were linked to the outbreak, ska distance was run on a merged skf file of the 45 outbreak isolates plus the 11 candidates. Results showed that then ten candidates that had perfectly matched the unique outbreak split kmers were all within 11 SNPs of an outbreak sample, while the candidate with only a 34% match to the unique outbreak split kmers was 53 SNPs from the closest outbreak sample. Ska type found consistent results, typing the ten closest samples as MLST ST2371, the novel ST identified in the SCBU outbreak samples, while the more distant candidate sample was typed as ST22. The ten samples were later confirmed as the same ten samples identified by Toleman *et al.* (Coll pers. comm.), illustrating the power of SKA to rapidly and accurately identify samples linked to an outbreak and to rule out links to other samples.

### Reconstructing the outbreak

To test the ability of SKA to reconstruct the phylogenetic relationships between isolates in an outbreak, it was compared with a traditional mapping approach similar to that used by Harris *et al.* and Toleman *et al.* in their epidemiological investigations. For transparency, the publicly available SNP pipeline (27) was used as the comparator. IQ-Tree was used to reconstruct phylogenetic trees from three alignments of 55 samples comprising the 45 original SCBU outbreak samples and the ten new samples identified by SKA:

1. A reference free alignment using ska align
2. An alignment created using SNP pipeline with the ST22 reference genome HO 5096 0412, the reference genome used by Harris *et al.* in their original investigation.
3. A gold standard alignment created using SNP pipeline with a complete reference genome of one of the samples from the outbreak identified by Harris *et al.*, which was subsequently sequenced using long-read PacBio sequencing (accession number GCA_001369415.1). All SNPs in the alignment were manually curated to ensure accuracy. This alignment was used to test the accuracy of alignments 1 and 2, which are more representative of real-life scenarios.

Table 4 provides an overview comparison of the ska alignment and the SNP pipeline alignment using the HO 5096 0412 reference, and figure 4 shows a comparison of the trees produced from these two alignments with the gold-standard tree created from alignment 3.

**Figure 4.**
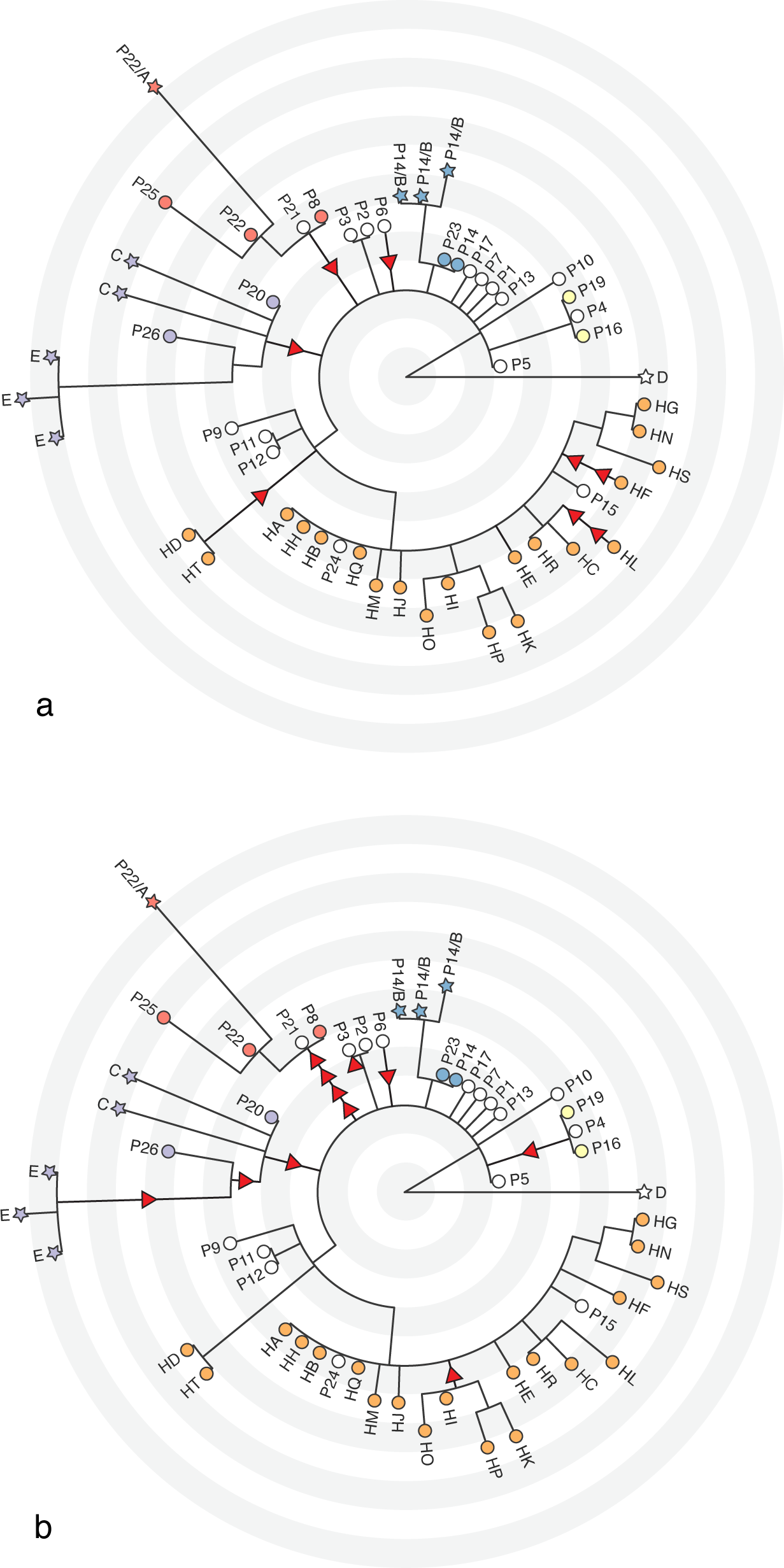
Accuracy of phylogenetic reconstructions of 45 isolates from the SCBU MRSA outbreak of Harris *et al.* (25) plus the ten samples from Toleman *et al.* (38) based on alignments created by ska align (a) and SNP pipeline mapped against a MRSA ST22 reference genome sequence (HO 5096 0412) (b). Both panels show how the trees reconstructed from these alignments compared with a gold standard tree created from a manually curated alignment based on a SNP pipeline mapping against a complete PacBio sequence of one of the outbreak samples (GCA_001369415). In all cases, trees were created from the alignments using IQ-Tree with a GTR+G+ASC model and the fast option. Red arrows indicate differences in branch length from the gold-standard tree. Arrows pointing towards the root indicate a reduction of a branch length by one SNP, while arrows pointing away from the root indicate an extension of a branch by one SNP. Multiple arrows on a branch indicate multiple SNP differences. The arrow below sample P3 in panel b is on the branch leading to sample P3, which was a zero-length branch in the gold-standard tree. Terminal node labels show samples names from Harris *et al.* (25) and Toleman *et al.* (38) Circles on terminal nodes indicate samples from Harris *et. al*. (25). while stars indicate samples from Toleman *et al.* (38). Their fill colour highlights samples from members of the same family (including multiple samples from the same individual). White fill indicates samples without family links. Each concentric circle of alternating white and grey indicates one SNP from the root, which was artificially placed on the branch leading to sample D.

**Table 4.** Comparison of SKA with SNP Pipeline, a typical mapping pipeline for analysis of bacterial genomic variation. These comparisons are based on analyses of 45 isolates from the SCBU MRSA outbreak of Harris *et al.* (25) plus the ten samples from Toleman *et al.* (38) with by ska align with the ‐v option to output an alignment of variable sites, and SNP pipeline under default options and using Bowtie2 for mapping. In both cases IQ-Tree was used to reconstruct a phylogeny from the alignment of variable sites under a GTR+ASC+G model and with the fast option. Note that the total disk space used could be reduced analyses if intermediate files were deleted during the analyses. *The maximum memory usage for SNP Pipeline was artificially reduced by restricting Picard MarkDuplicates and GATK RealignerTargetCreator and IndelRealigner to 5Gb RAM.

|  | SKA | SNP Pipeline | Fold difference |
| --- | --- | --- | --- |
| **Total topological differences to gold standard tree** | 0 | 0 | - |
| **False positive SNPs** | 0 | 5 | - |
| **False negative SNPs** | 8 | 7 | 0.875 |
| **Total CPU time (m)** | 114 | 1,913 | 16.78 |
| **Total CPU time (h)** | 1.9 | 31.9 |  |
| **Maximum memory usage (Mb)** | 607 | 4,479* | 7.38 |
| **Maximum file size (Mb)** | 31 | 2,450 | 79.03 |
| **Total disk space used (Mb)** | 1,661 | 116,746 | 70.29 |

The trees created from each alignment and associated metadata including sample accession numbers are available at, https://microreact.org/project/SKASaureusSKAalign, https://microreact.org/project/SKASaureusSNPPipeline and https://microreact.org/project/SKASaureusGold, respectively. In all cases the topologies of the trees were consistent, however branch lengths differed in correspondence to the number of false positive and false negative SNPs for each method. In summary, SKA failed to identify eight variant sites, while SNP Pipeline missed seven and erroneously called SNPs at another five sites. Four of the false negatives in the SKA analysis were the result of SNPs in repeats that were longer than the total split kmer size. The other four were the result of two cases where variant sites were within a kmer length of each other. Such an occurrence is usually uncommon in datasets with such low levels of variation, but serves to illustrate the limitations of the method. Three of the false negatives in the SNP Pipeline analysis were in a region that was not present in the reference genome. Three more were removed by the pipeline default filters as there were more than three SNPs in a region of 1000bp in some samples. The final false negative was in a region duplicated in the outbreak samples relative to the reference, and corresponded to one of the false negatives of SKA. Four of the false positives called by SNP Pipeline were mis-alignments around indels relative to the reference, while the fifth was a heterogeneous call in a repeat region of varying length. It is important to note that even the gold standard alignment contained erroneous SNP calls prior to manual curation and phylogenetic reconstruction. In total nine false positives were removed. Five of these were mis-alignments around indels relative to the reference and another four were heterogeneous calls in repeat regions.

Computational comparisons between the alignment methods showed much greater disparity (Table 4). SNP Pipeline required nearly 17 times more CPU time than SKA and over 70 times more disk space. Memory usage was also 7 time greater and would have been higher had the Picard MarkDuplicates and GATK RealignerTargetCreator and IndelRealigner parts of the pipeline not been restricted to 5Gb RAM. Furthermore, the most CPU intensive part of the SKA analysis is the creation of a merged skf file from the raw sequencing reads.

Alignment from this file took only 20s and 208Mb RAM. The reference-free approach means that there would not be a need to remap to a different reference, but other analyses from the file, including mapping to a reference genome or distance calculation, would be similarly fast. Remapping to a different reference using a mapping approach such as SNP Pipeline, on the other hand, would require the same computational resources as for the first reference: in this example, that involved over 30 CPU hours, 5Gb RAM (or more if run in parallel) and over 100Gb disk space.

## Discussion

Genomics is becoming increasingly used in epidemiological investigations of bacterial pathogen outbreaks and to understand their transmission dynamics. However, analysis of genomic data often requires the skills of a dedicated bioinformatician and large compute resources. Results can depend on choices which need to be made during those analyses, such as the reference used for read mapping or the assembler used to create contigs. Thus, a level of understanding of genomics is required to both analyse the data and interpret the results.

Here I have introduced a new approach to genomic analysis of highly-related, small, haploid genomes that lends itself well to epidemiological uses. In tests using published simulated and real-life data, SKA compared well to the best of the currently-used methods. However, SKA has a number of advantages over these methods. It is not dependent upon other bioinformatics software, is computationally efficient and produces small intermediate files, allowing analyses to be carried out on a typical laptop or desktop computer very quickly. Importantly, SKA allows genomic analysis that requires few, or no choices to be made by the user, including reference-free analyses that are not restrained by the need for a pre-defined core genome or closely-related reference sequence. A simple automated pipeline could convert raw sequence reads to skf files directly after sequencing, merge them using ska merge, cluster using ska distance, align each cluster using ska align and reconstruct trees using the algorithm of choice, all without even needing to know the species being sequenced. If an outbreak is identified, SKA includes methods identify outbreak-defining split kmers that can be used to quickly scan large samples sets for related samples. The ability to annotate split kmers onto reference genome sequences allows the biological consequences of variants identified during these analyses to be assessed.

The main limitations of using split kmers are similar to all other kmer-based methods. In particular, their inability to identify SNPs in repeated split kmers or in cases where variant sites are within a kmer length of one another or of an indel can lead to loss of some signal. Fortunately, such cases are usually rare in outbreak scenarios. However, future improvements could allow post-processing of split kmers to identify these occurrences without impacting greatly on speed or efficiency. Traditional methods also have limitations. Core genome methods have reduced sensitivity due to the exclusion of large parts of the genome from the analysis. Mapping-based approaches suffer from more opaque biases. The distance of the reference genome from the samples of interest, for example, leads to reference-sample-specific mapping biases. SNPs in regions missing from the reference will be excluded from the analysis and indels or repeats relative to the reference can lead to false positive SNP calls, even when the reference is very closely-related to the samples being analysed. Thus, comparison between analyses using different reference genomes is not ideal, although often is necessary. Furthermore, the reliance of the majority of mapping pipelines on many independent software tools makes versioning extremely tricky. For example, SNP Pipeline requires the installation of ten external dependencies. To ensure reproducibility when analyzing data in different locations all dependencies should be of the same version, something that is far from straightforward to manage. Biases caused by this type of variation in pipelines are usually completely ignored.

In summary, SKA is a powerful new toolkit for the genomic epidemiologist with the potential to speed up outbreak detection and investigation from short read sequencing.

## Acknowledgements

I would like to thank John Lees for help with Kendall Colijn metric analysis, Ben Taylor for testing SKA on various datasets, and Francesc Coll and Michelle Toleman for help accessing and linking their data to appropriate metadata. I would also like to thank all at the Centre for Genomic Pathogen Surveillance and the Wellcome Sanger Institute who provided feedback and the Pathogen Informatics team at the Wellcome Sanger Institute for informatics support. This work was funded by Wellcome grant number 098051.

